# Loci, genes, and gene networks associated with life history variation in a model ecological organism, *Daphnia pulex* (complex)

**DOI:** 10.1101/436733

**Authors:** Jacob W. Malcom, Thomas E. Juenger, Mathew A. Leibold

## Abstract

**Background:** Identifying the molecular basis of heritable variation provides insight into the underlying mechanisms generating phenotypic variation and the evolutionary history of organismal traits. Life history trait variation is of central importance to ecological and evolutionary dynamics, and contemporary genomic tools permit studies of the basis of this variation in non-genetic model organisms. We used high density genotyping, RNA-Seq gene expression assays, and detailed phenotyping of fourteen ecologically important life history traits in a wild-caught panel of 32 *Daphnia pulex* clones to explore the molecular basis of trait variation in a model ecological species.

**Results:** We found extensive phenotypic and a range of heritable genetic variation (~0 < H^2^ < 0.44) in the panel, and accordingly identify 75-261 genes—organized in 3-6 coexpression modules—associated with genetic variation in each trait. The trait-related coexpression modules possess well-supported promoter motifs, and in conjunction with marker variation at trans- loci, suggest a relatively small number of important expression regulators. We further identify a candidate genetic network with SNPs in eight known transcriptional regulators, and dozens of differentially expressed genes, associated with life history variation. The gene-trait associations include numerous un-annotated genes, but also support several a priori hypotheses, including an ecdysone-induced protein and several Gene Ontology pathways.

**Conclusion:** The genetic and gene expression architecture of *Daphnia* life history traits is complex, and our results provide numerous candidate loci, genes, and coexpression modules to be tested as the molecular mechanisms that underlie *Daphnia* eco-evolutionary dynamics.

## INTRODUCTION

Determining the relationship between genetic, trait, and ecological variation is a key goal of contemporary biological research. Investigating the relationship between genotype and phenotype—including genotype x environment interactions—is the domain of genetics, which rarely extends to higher levels of organization such as populations and communities. A complement at higher levels of biological organization is the field of trait-based ecology, which focuses on the relationship between phenotype and ecological processes such as population dynamics and community assembly [1–4]. A key step forward is to solidify these cross-hierarchy links in ecologically well-studied settings, but this has not yet been accomplished. The main reasons for this shortcoming are two-fold: we know very little about the ecology, and the ecological context of evolution, of most model genetic organisms; and genetic resources are sparse for model ecological organisms [5–8]. Therefore, an ideal system with which to approach such a problem is one for which we possess both extensive genetic resources and ecological knowledge.

With the recent genome sequencing of the waterflea, *Daphnia pulex* (hereafter *Daphnia*), we can integrate information across levels of organization, from genetic sequence to ecologically important traits for an organism that is generally considered a keystone species [9, 10]. The *Daphnia* genome is approximately 227MB in length and is characterized by numerous tandem duplications with rapid divergence of expression patterns [11]. A century of ecological research has shown that *Daphnia* are central players in aquatic communities by acting as a key link between producers and carnivores; by controlling plant biomass and production; and by their effects on nutrient cycling in lakes (see [12] for a brief review). Arguably, we know the factors describing the requirement and impact niches [13] of *Daphnia* as well as, if not better than, any other species. Furthermore, we know that rapid adaptation in *Daphnia* can have dramatic effects on ecological dynamics [14–17]. The nexus of ecological and evolutionary knowledge with genomic tools for a single species enables linking the levels of biological organization in a way not possible for classical model genetic species and other model ecological species.

Several life history traits are central to shaping ecological and evolutionary dynamics, both in *Daphnia* and more generally. To provide context, we briefly review the general relevance of size, growth, and fecundity to ecological variation, including the specific connections to *Daphnia* biology. Next, we set out several a priori hypotheses concerning the genes and biological pathways we might expect, based on the literature, to be associated with variation in the life history traits examined. Finally, we define the goals of the present study— which are largely descriptive—before moving on to the results.

### The ecological importance of life history traits

Life history traits are among the ecologically most-important for a wide array of taxa. Few traits are thought to affect ecological dynamics more than body size [18, 19]: metabolism scales with body size and is predictive of macroecological patterns [20, 21]; body size limits the upper size of food an individual can consume (e.g., [22]), and body size is intimately tied to predation susceptibility [23]. Size is also strongly correlated with the number and size of offspring produced [24], which is a vital aspect of fitness and population growth. Individuals who tend to grow faster are larger as adults and tend to require more resources than individuals who are smaller. As a result, the nutritional requirements are greater, and the impacts on resource availability are greater, for larger individuals [25].

These (and many other) trait:ecology mappings have been examined in *Daphnia*. Patterns of cladoceran body size, and *Daphnia* in particular, formed the empirical basis for Brooks and Dodson’s influential size efficiency hypothesis [26]. They argued that larger species such as *Daphnia* are better competitors and should dominate any given community except in the face of vertebrate predators. While true in some cases, there are many caveats to the general pattern [27], including the fact that competitive outcomes with respect to body size are conditional on food quantity and quality: larger *Daphnia* have an advantage when algae is abundant and high-quality, but smaller organisms or individuals gain the upper hand when algae density or quality is low [28, 29].

While size confers certain competitive advantages and disadvantages, it also shapes predation risk. Large *Daphnia* are more visible and therefore more susceptible to predation by vertebrate predators [30–32]. Rapid evolution—within a single season, and spanning just a few generations—of body size in response to seasonal changes in fish predation regimes has been demonstrated in the species [33, 34]. In contrast to the interaction of size and vertebrate predators, small *Daphnia* are susceptible to predation by invertebrates, including the well-studied effects of the midge larvae (*Chaoborus* spp.) and copepods, which are unable to effectively handle larger prey [35–38]. Furthermore, vertebrate and invertebrate predator regimes are not independent of one-another [37, 39], nor are they independent of other ecological processes such as competitive interactions [40]. Body size and its associated traits are thus key traits that mediate the relative susceptibility of *Daphnia* to different predators, and to be efficient resource exploiters in different habitats.

As competition and predation impose selection on *Daphnia* body size, correlated traits are also affected. For example, because reproductive output is often strongly correlated with maternal size, we expect that selection regimes favoring smaller *Daphnia* will also results in fewer and/or smaller offspring and affects influences ecological dynamics [33]. The size-efficiency hypothesis suggests that larger offspring have greater competitive ability when food is scarce, but more, smaller offspring provide a numerical advantage when food is abundant; Tessier and Goulden [29] refined the hypothesis to state that larger individuals are at an advantage when food availability fluctuates extensively. Larger offspring tend to have higher growth rates, however, which requires higher nutrient concentrations to support growth (in particular, phosphate; [41, 42]). The nutrient environment in which *Daphnia* grow and reproduce is shaped by abiotic and biotic factors, e.g., shading alters algal stoichiometry to increase the amount of phosphorus relative to carbon, which interacts with genotype to alter growth rate and body size [43–47].

### Expectations

While the genomic resources for *Daphnia* are a recent development, we can make several predictions about the genes and pathways expected to be associated with life history trait variation. Genome-wide expression and genotyping studies facilitate discovery of novel gene-trait associations [48], but post-hoc explanations of patterns are weaker than tests of a priori hypotheses. Decades of classic molecular, reverse-, and forward-genetic studies provide guidance as to the pathways and genes expected to be related to trait variation; before we analyzed any SNP-trait or expression-trait data for *Daphnia*, we therefore searched the existing literature for candidate genes and pathways that might allow us to make a priori hypotheses. Recovering expression or genotype variation in these pathways, related to trait variation, provides additional support for the statistical inference.

Insulin and insulin-like signaling pathways are commonly related to size- and growth-related traits in numerous model species [49–52]. Ecdysone-activated proteins, which regulate arthropod molting and were the top candidates in experimental evolution of fly body size [53], are found in *Daphnia* (e.g., nucleolar protein c7b, which is a homolog of the *Drosophila* gene *mustard*). FOXO, a forkhead transcription factor, is central to insulin signaling and stress response, the latter of which was found to be a top body size-related pathway using flies from the *Drosophila* Genetic Reference Panel (DGRP; [54, 55]. Neuronal control of size—through related behaviors such as movement patterns and feeding rate—has been demonstrated, for example with *C. elegans* (egl-4, a cGMP-dependent kinase; [51]) and *Drosophila* (short neuropeptide F, a protein precursor thought to be central to chemosensation; [56]). Although some of these examples are very specific, they provide guidance on what to expect when exploring the genomics of life history variation in a newly developing model species such as *Daphnia*.

Less information is available about the genes and pathways underlying variation in the number of offspring or time between broods (clutches). The Gene Ontology categories for genes best-correlated to absolute fitness in flies from the *Drosophila* Genetic Reference Panel include proteolysis, signal transduction, and defense/immune response [57]. There are numerous vitellogenin-like genes in the *Daphnia* genome, and contaminant stressors have been shown to affect the expression of vitellogenin genes and the levels of the vitellogenin antagonist, juvenile hormone in *Daphnia* magna [58]. The copy number of yolk protein genes (*yp-1*, *yp-2*, and *yp-3*) is positively correlated with egg production in *Drosophila* [59], but there are no clear *Daphnia* homologs (the closest BLAST hits are to a pancreatic triacylglycerol lipase). Of note, the yp genes share sequence similarity with vertebrate lipases, which is thought to underlie the ability to store various steroids used in developmental signaling [60]. We expect to find support for some or all of these genes, plus new candidates, by association with variation in *Daphnia* fecundity traits.

Long gene lists are often of less utility than general terms associated with the function of groups of genes for understanding the biology of variation in quantitative traits. After considering a combination of the literature and basic biology, we developed a matrix of biological terms, guided by the Gene Ontology [61] and KEGG [62] frameworks, expected to be enriched for groups of the growth and fecundity traits (Table 1). These relationships are straight-forward, e.g., we anticipate enrichment for metabolism and genetic information processing groups of terms because of the need to convert materials into biomass, and several signaling and cell cycle-related terms because of their role in development and growth.

**Table 1.**
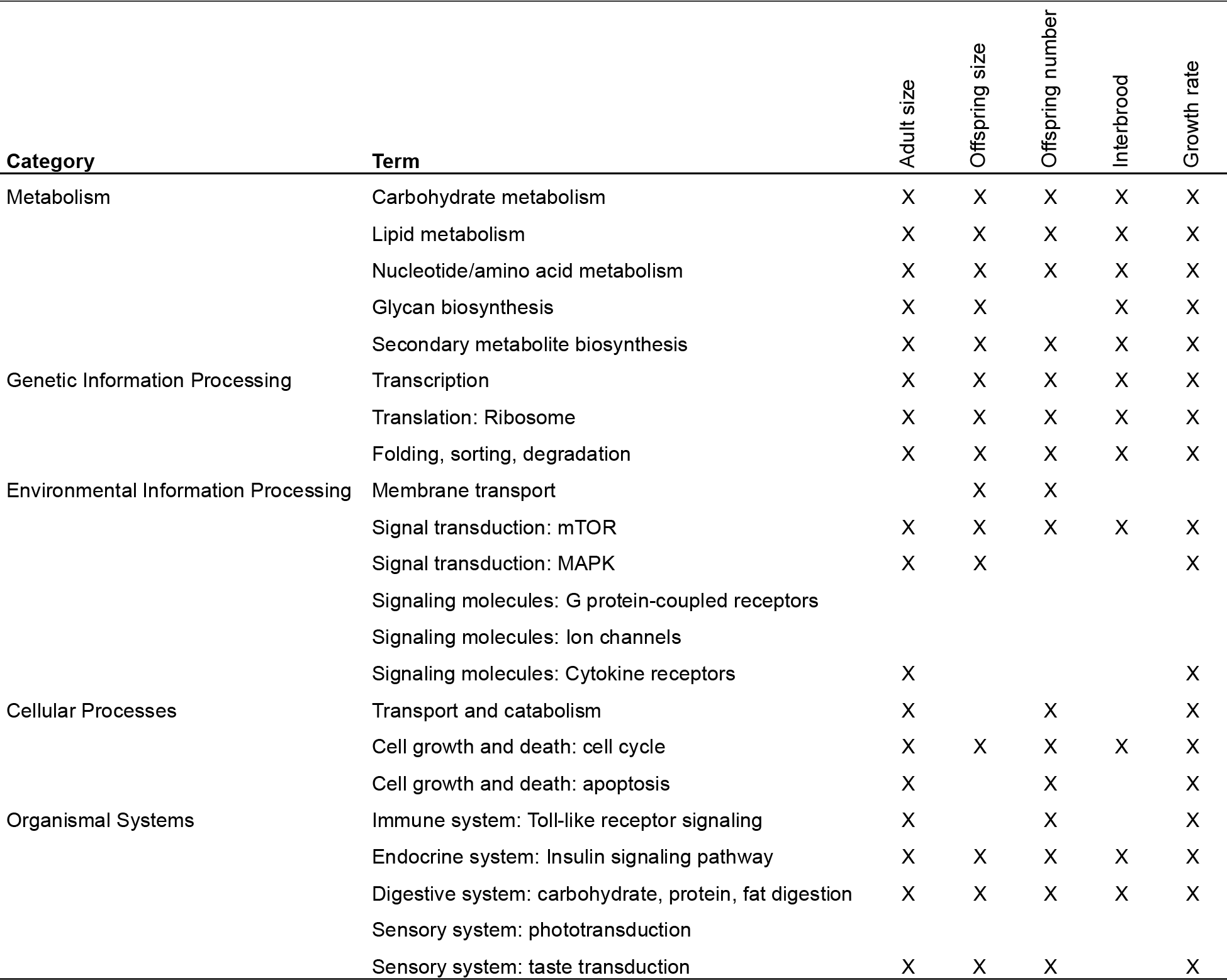
Expected trait-pathway relationships for groups of *Daphnia* growth and fecundity traits. The categories and terms are derived from KEGG and Gene Ontology.

### Present goal

Given the general ecological importance of size, growth, and fecundity; the well-established role variation in these traits plays in shaping *Daphnia* ecological and evolutionary dynamics; and a desire to understand the connections across the genetic, trait, and ecological levels of organization, we investigated the molecular basis of life history variation in a panel of wild-collected *Daphnia*. We quantified variation in fourteen growth and fecundity traits, used RNA-Seq to quantify constitutive gene expression variation across the panel, and genotyped each clone at an average of ~3 million loci. We then integrated this data to provide estimates of the relationships between genetic, expression, and phenotypic variation. The results indicate many genes organized in a relatively small number of coexpressed gene modules are associated with variation in these traits, provide novel hypotheses to be explored as the field of ecological genomics expands, and establish several candidates for genes underlying the interface of eco-evolutionary dynamics.

## RESULTS

### Phenotypic variation

There was substantial variation across the panel of 32 clones for all 14 traits that we measured. First, the difference between the smallest and largest clones was approximately four-fold for juvenile mass and three-fold for adult mass (Figures 1A-C). The differences were smaller for *Daphnia* length and depth (Figures 1D-E), but of similar magnitude for growth rate (Figure 1F). The variation in body sizes translated to substantial variation in the number of offspring, with up to a 9-fold difference in brood size (Figures 2A-C). In addition, there was up to a three-fold difference in interbrood period (Figures 2D-F). Clone-wise summary statistics are provided in Supplemental Information (SI) Table 1.

**Figure 1.**
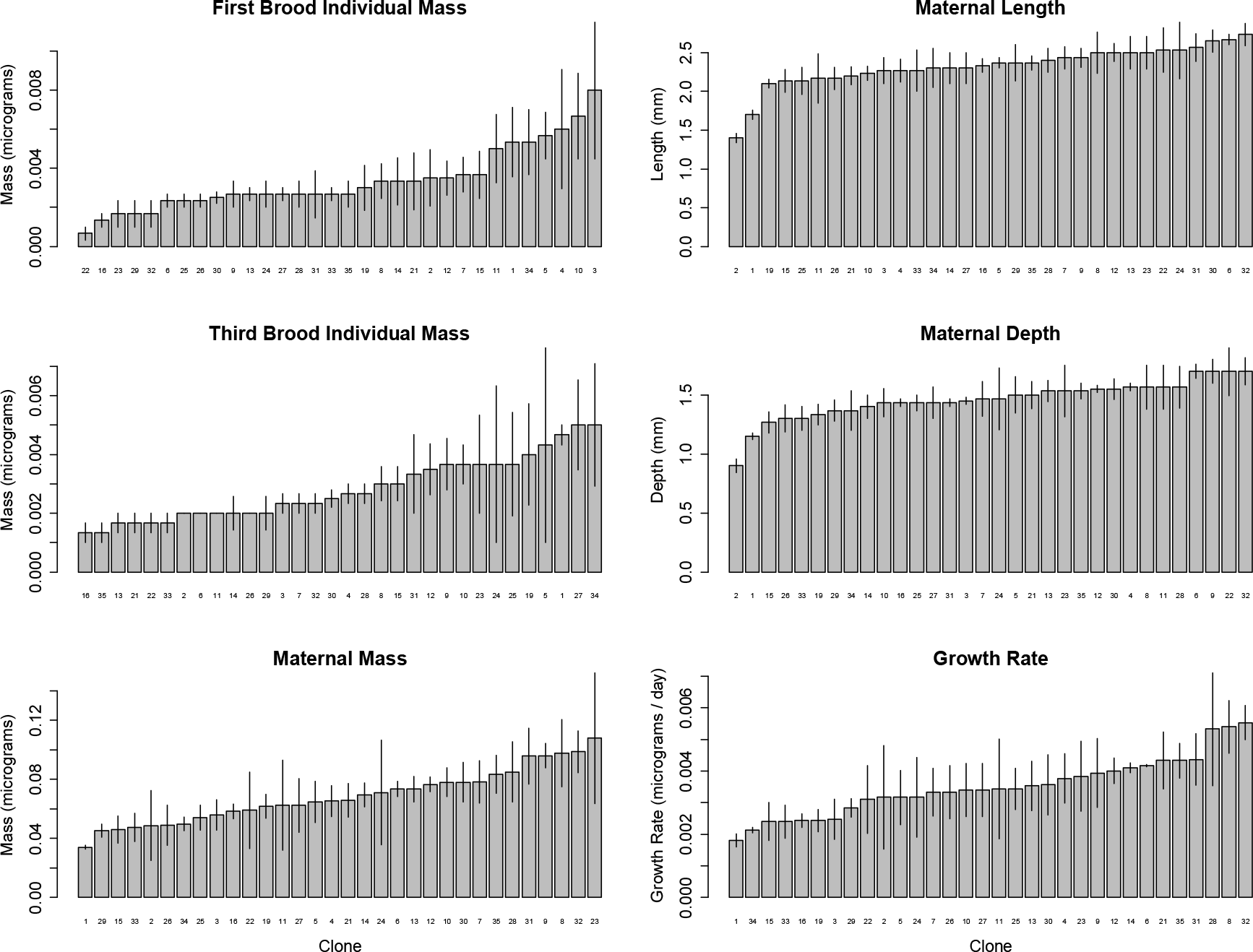
Panel variation of *Daphnia* size traits. First and third brood individual mass is the mass of individual offspring, maternal size characteristics are measured at the third brood, and growth rate is calculated from the maternal mass, first brood individual mass, and time from birth to third brood (see Figure 2). Error bars are 95% confidence intervals, and the clone-wise summary data is provided in SI Table 1.

**Figure 2.**
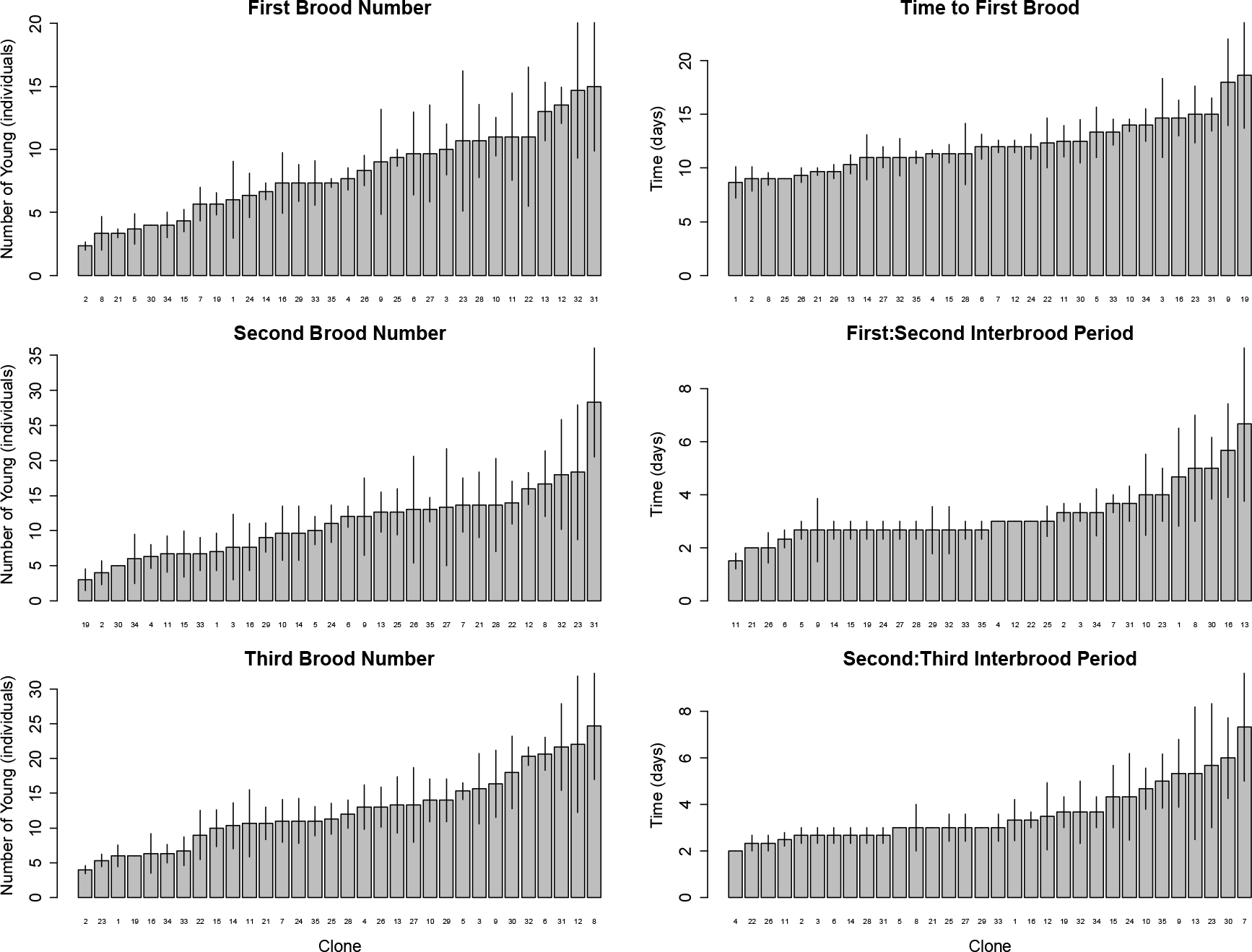
Panel variation in the number and timing of offspring. Error bars are 95% confidence intervals, and the clone-wise summary data is provided in SI Table 1.

Broad-sense heritability varied from ~0 to 0.44 for the life history traits, but some evolvabilities (i.e., mean-scaled genetic variance) were particularly high although heritability was relatively low (e.g., number and mass of first-brood offspring; Table 2). Phenotypic correlations (Figure 3, above-diagonal) were generally weaker than genetic correlations (Figure 3, below diagonal), and each tended to be in the directions expected given prior results with *Daphnia* and life history research in general. Adult mass was positively correlated with external measurements, the number of offspring per brood, time to first reproduction and the interbrood period. Adult mass was negatively correlated with the mass of offspring in the first brood; larger females invested in more offspring rather than allocating resources to larger offspring. Although the relationship was weaker, the number of offspring in a brood was negatively correlated with the size of those offspring. One prominent exception to the expected correlations was the negative relationship between offspring size and growth rate: smaller neonates grew faster than larger neonates. Interestingly, environmental variance appears to canalize some correlations, such as the weakly negative genetic correlations between adult body size (depth and length, −0.13 > *r* > −0.15) and mass of third-brood offspring, to strongly negative (*r* ~ −0.29).

**Table 2.** Quantitative genetics of growth and fecundity traits in the *Daphnia* panel. H^2^ = broad-sense heritability; CVg = genotypic coefficient of variation; Vg,p = genotypic and phenotypic variances; mean = mean trait value across all replicates of all clones.

| Stage | Trait | mean | $V_g$ | s.d.($V_g$) | $V_R$ | s.d.($V_R$) | $V_p$ | $H^2$ | LCL($H^2$) | UCL($H^2$) | $I_g$ |
| --- | --- | --- | --- | --- | --- | --- | --- | --- | --- | --- | --- |
| Offspring | # (brood 1) | 8.089 | 4.60E+00 | 2.15E+00 | 2.14E+01 | 4.62E+00 | 2.60E+01 | 0.18 | 0.15 | 0.20 | 7.04 |
|  | mass (br. 1, mg) | 0.003 | 1.03E-06 | 1.01E-03 | 4.97E-06 | 2.23E-03 | 6.00E-06 | 0.17 | 0.14 | 0.20 | 9.18 |
|  | # (br. 2) | 11.156 | 1.51E-08 | 1.23E-04 | 6.61E+01 | 8.13E+00 | 6.61E+01 | 0.00 | -0.03 | 0.03 | 0.00 |
|  | # (br. 3) | 12.653 | 1.31E+01 | 3.62E+00 | 4.48E+01 | 6.69E+00 | 5.79E+01 | 0.23 | 0.20 | 0.25 | 8.20 |
|  | mass (br. 3) | 0.003 | 0.00E+00 | 0.00E+00 | 3.67E-06 | 1.92E-03 | 3.67E-06 | 0.00 | -0.03 | 0.03 | 0.00 |
| Adult | mass (mg) | 0.068 | 2.67E-14 | 1.64E-07 | 9.04E-04 | 3.01E-02 | 9.04E-04 | 0.00 | -0.03 | 0.03 | 0.00 |
|  | length (mm) | 2.322 | 3.59E-02 | 1.89E-01 | 1.00E-01 | 3.17E-01 | 1.36E-01 | 0.26 | 0.25 | 0.28 | 0.67 |
|  | depth (mm) | 1.459 | 1.20E-02 | 1.10E-01 | 4.63E-02 | 2.15E-01 | 5.83E-02 | 0.21 | 0.19 | 0.23 | 0.56 |
|  | tailspine (mm) | 0.631 | 9.51E-03 | 9.75E-02 | 1.19E-02 | 1.09E-01 | 2.14E-02 | 0.44 | 0.43 | 0.46 | 2.39 |
|  | time to br. 1 (days) | 12.146 | 2.50E+00 | 1.58E+00 | 1.07E+01 | 3.27E+00 | 1.32E+01 | 0.19 | 0.16 | 0.22 | 1.70 |
|  | time br. 1 to br. 2 (d) | 3.255 | 4.59E-10 | 2.14E-05 | 3.12E+00 | 1.77E+00 | 3.12E+00 | 0.00 | -0.04 | 0.04 | 0.00 |
|  | time br. 2 to br. 3 (d) | 3.604 | 2.78E-10 | 1.67E-05 | 4.18E+00 | 2.05E+00 | 4.18E+00 | 0.00 | -0.03 | 0.03 | 0.00 |
|  | growth rate (mg d<sup>-1</sup>) | 0.004 | 1.49E-16 | 1.22E-08 | 2.33E-06 | 1.53E-03 | 2.33E-06 | 0.00 | -0.02 | 0.02 | 0.00 |

**Figure 3.**
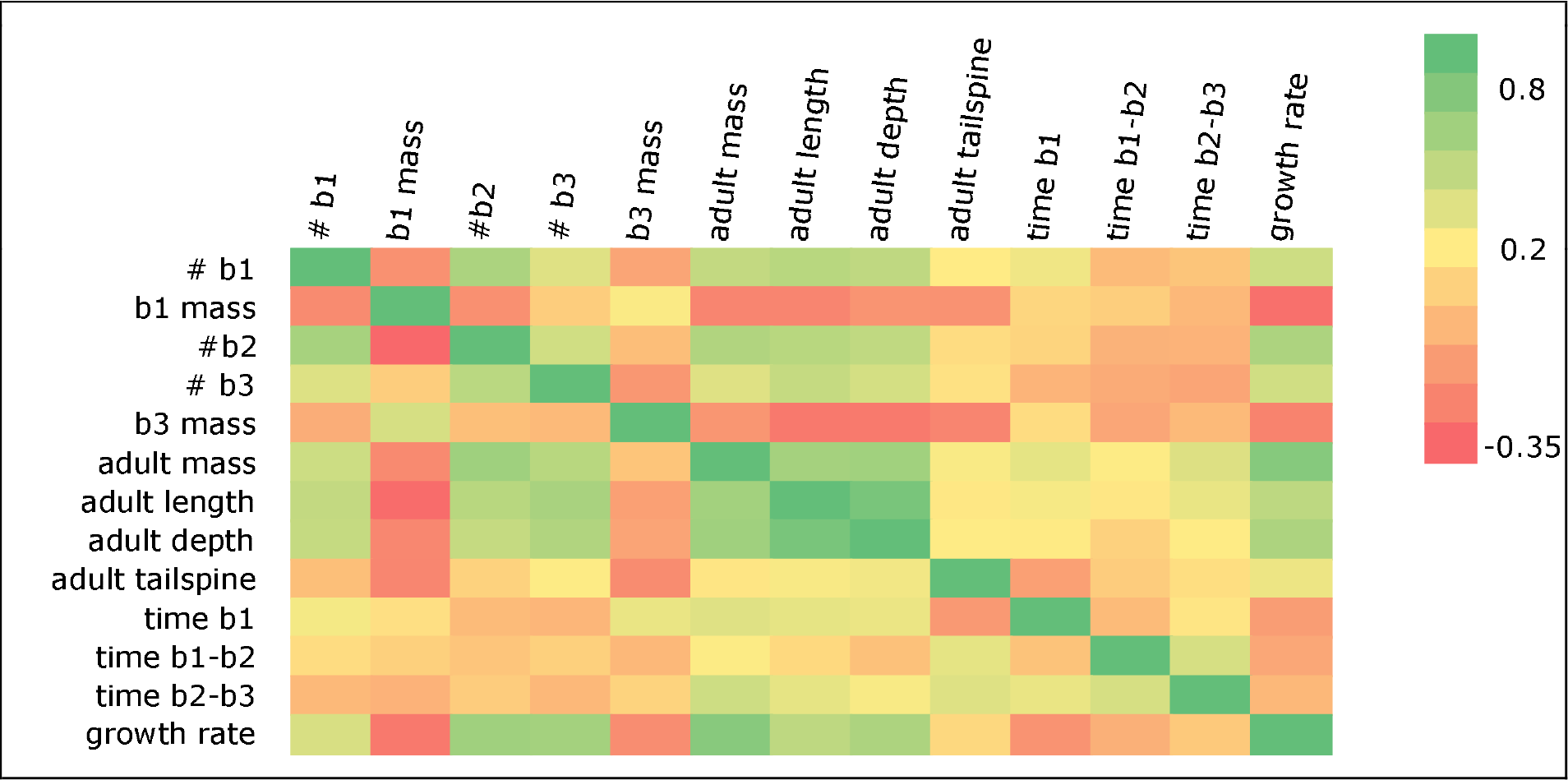
Genetic (below diagonal) and phenotypic (above diagonal) inter-trait correlations among growth and fecundity traits. “bN” is the brood number of interest. Note that phenotypic correlations tend to be weaker than genetic correlations, but there are exceptions such as the stronger phenotypic correlation between b3 mass and several other traits.

### Genetic variation

Sequencing from 2b-RAD and RNA-Seq libraries resulted in an average 2.2 × 10^6^ raw reads per clone, which, when aligned to the *Daphnia* reference, resulted in 5.5 × 10^6^ genotyped base positions across the population and an average of 4.66 × 10^6^ loci genotyped per clone (s.d. = 1.2 × 10^6^). There were 6338 loci polymorphic in the panel and typed in >75% of clones; heterozygosity rate was low, with a mean across clones of 0.14%. Mean indel and substitution rates estimated during read mapping were 2.7 × 10^−4^ and 0.022, respectively. We observed few patterns of genome-wide genetic variation; for example, mean nucleotide diversity in 50kb sliding windows was relatively constant (Figure 4). Population structure is relatively low in the panel, and linkage disequilibrium appeared to decline relatively quickly, to background levels by about 200bp (see SI Text 1).

**Figure 4.**
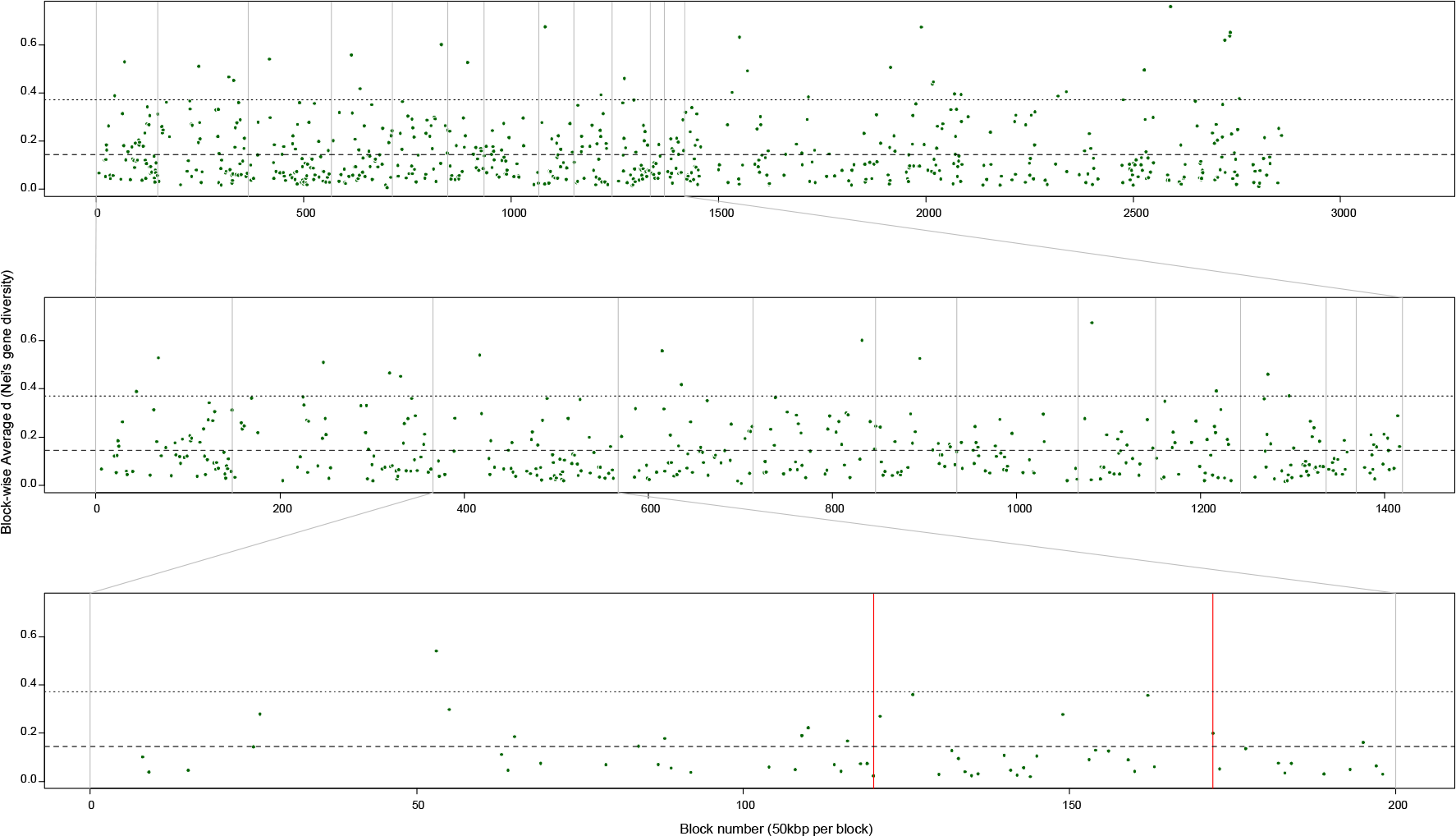
Nucleotide diversity (Nei’s d) across the genome (A), among scaffolds anchored to linkage groups (B), and within linkage group 3 (C). Light gray vertical lines demarcate linkage group boundaries, and the vertical red lines in panel C mark the window containing LDH genes, which is a common marker for clone identification in ecological studies. Horizontal dashed lines are the mean diversity values and horizontal dotted lines mark the 95th percentile of estimated diversity values. Nucleotide diversity was calculated either in 50-kb non-overlapping windows or over the entirety of a scaffold (if less than 50kb), and only if there were ≥ 3 polymorphic, typed loci in the window/scaffold. Ordering follows the marker order given on the *Daphnia* Genomics Consortium web portal; scaffolds with markers whose order was ambiguous or conflicting relative to the genome assembly were considered unanchored and move to the unanchored section of panel A.

### Gene expression variation

As expected given the wide variation in size, growth, and fecundity traits, we uncovered substantial variation in constitutive gene expression across the 32 clones. Although > 30,000 *Daphnia* genome features (i.e., genes) possessed at least one mapped read, subsequent analyses are based on a subset of 15,600 genes with mean expression of 5 reads per million mapped. Approximately 45% (n = 6434) of genes were differentially expressed (DEGs) at p < 0.05 after Benjamini-Hochberg FDR control at a 1% rate, using a generalized linear model with quasipoisson errors (see METHODS). DEGs possess a significantly higher broad-sense heritability (mean H^2^ = 0.657) compared to all expressed genes (mean H^2^ = 0.508, *p* < 2.2e-16; SI Figure 1). The degree of expression differences varied from 1.6- to 7000-fold, due only to genetic differences between clones in a common garden environment.

Global gene expression was highly modular, with 24-27 distinct coexpression modules recovered across a wide range of parameter conditions. The genes within each module were associated with 1-17 novel promoter motifs (SI Table 2) based on word enrichment in the sequence 1000bp upstream and 200bp downstream of the transcription start site, as determined with XXmotif (see METHODS). Each module was enriched for 6-97 Gene Ontology (GO) terms, ranging from generic “biological process” to more detailed “hydrolase activity” (SI Table 3).

**Table 3.**
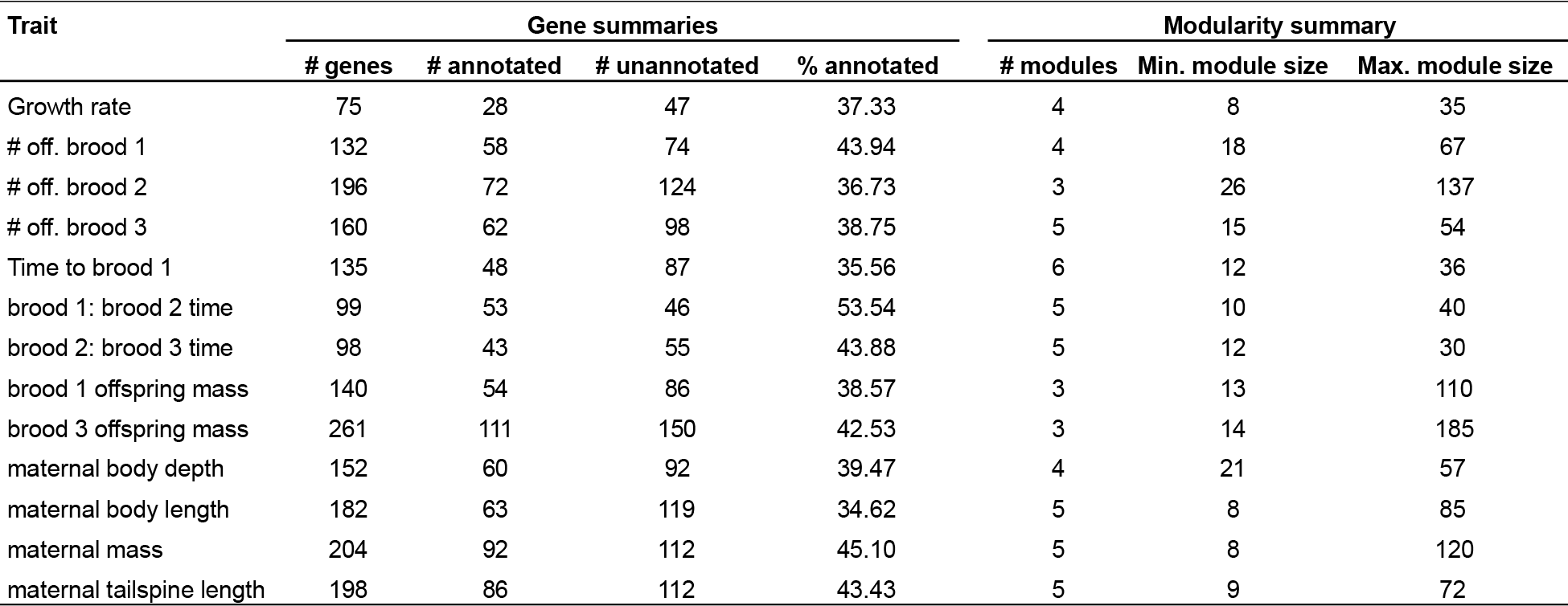
Summary statistics for differentially expressed genes (DEGs) and coexpression modules associated with variation in the fourteen *Daphnia* growth and fecundity traits. The specific genes and module memberships for each trait are provided in SI Table 5.

### Linking genotypic, expression, and trait variation

We identified 14 SNPs (nominal *p* < 1e^−5^) and an average of 156 DEGs (*p* < 0.05 and FDR control at 1%) whose variation was tightly correlated with variation in each of the fourteen growth and fecundity traits (Table 3, SI Table 4). Two gustatory receptors (hxAUG25s1441g78t1, hxJGI_V11_235732) were differentially expressed and associated with second and third brood sizes, but no insulin receptor or peptide-encoding genes were associated with growth and fecundity traits. One SNP (scaffold 17:1051920) was located in an ecdysone-induced protein gene, and one differentially expressed ecdysone receptor (hxAUG26res30g96t1, 20-hydroxy-ecdysone receptor 20e) was associated with variation in offspring mass and adult tailspine length. Several lipases are differentially expressed and strongly correlated with number of offspring, time between broods, and adult tailspine length. There are currently 26 vitellogenin-associated genes annotated in the *Daphnia* genome; expression of only one vitellogenin precursor, hxJGI_V11_307854, was associated with variation in adult body length, but not brood size, offspring size, or time between broods.

Genes associated with each life history trait clustered into 3-6 modules of coexpressed DEGs per trait (Table 3, SI Figure 2), drawn from an average of 4.3 global expression modules (range 2-8; SI Figure 3). There were substantial differences in the architecture of trait-specific coexpression networks. For example, while there is a similar number of genes associated with adult mass (Figure 5A) and third brood offspring mass (Figure 5B), the adult mass network has more “hub” genes (i.e., with high betweenness-centrality, a metric of the node’s importance in connecting the network) than the offspring mass network. The adult mass gene with the highest betweenness centrality is *Partner of bursicon*, a precursor for a neurohormone involved in molting [63]; the two hub genes of the offspring mass network encode ERK-a and Rab-32 proteins, members of the MAPK/ERK signaling pathway [64].

**Figure 5.**
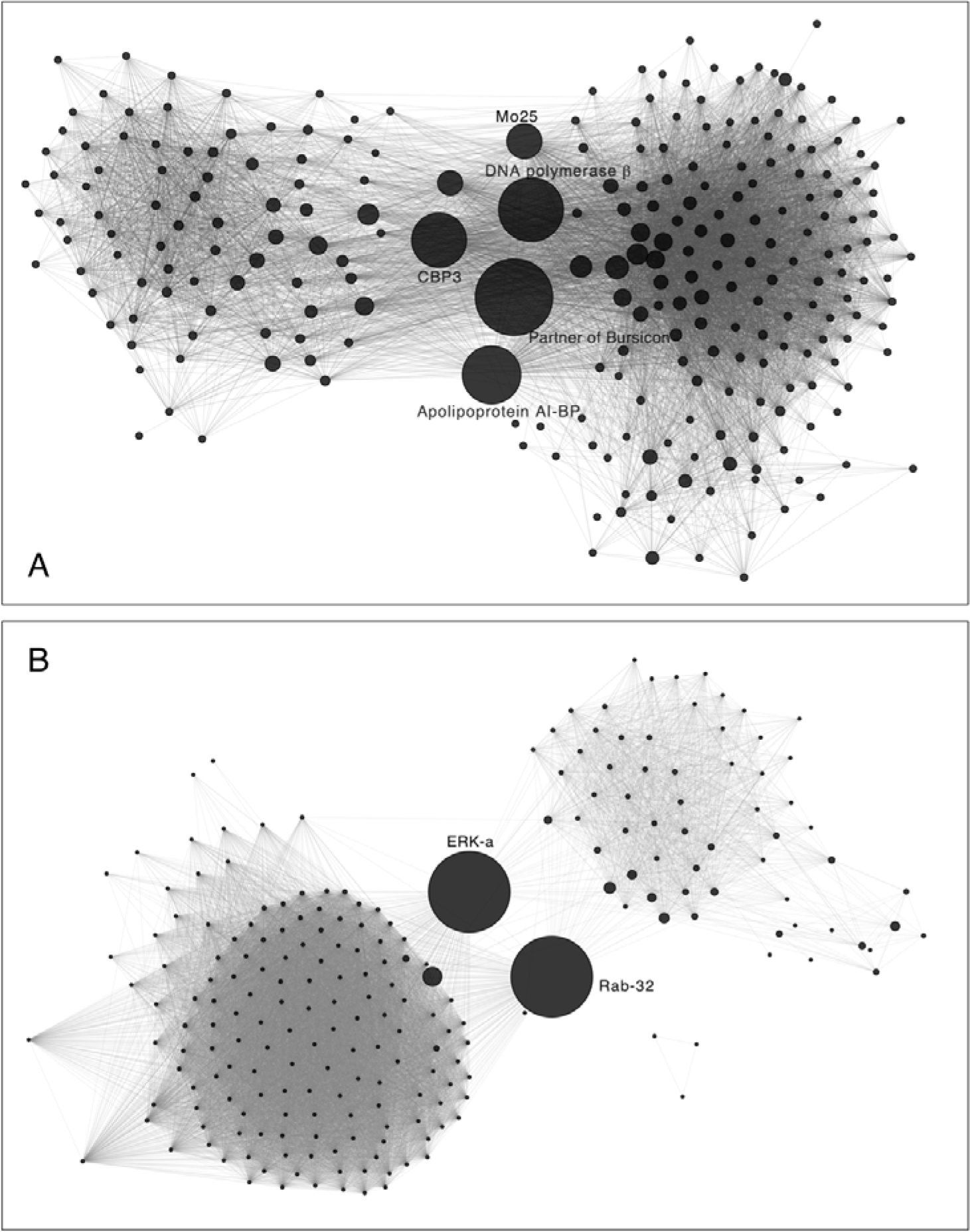
Coexpression networks associated with variation in adult mass (A) and third-brood offspring mass (B). Node size is proportional to the betweenness-centrality of the gene; larger nodes connect more groups of coexpressed genes. Select genes with high betweenness-centrality are labeled and discussed in the text.

Although 60% of genes associated with *Daphnia* growth and fecundity variation possess no functional annotation, we identified between seven and 79 GO terms enriched for each growth and fecundity related trait (SI Table 5). Across all fourteen traits, GO terms associated with gene expression regulation, protein transport, metabolism, cell proliferation, and signaling were among the top enriched Biological Process (BP) groups (Figure 6A). Correspondingly, nucleic acid binding, transporter activity, macromolecule binding, and receptor activity were among the groups of Molecular Function terms enriched (SI Figure 4). Traits characteristic of adult *Daphnia* (i.e., size and growth rate characteristics) were enriched for BP terms related to protein transport, cell cycle and proliferation, and growth (Figure 6B), while offspring-related traits were enriched for gene expression regulation, developmental processes, signaling, and cell death-associated BP terms (Figure 6C).

**Figure 6.**
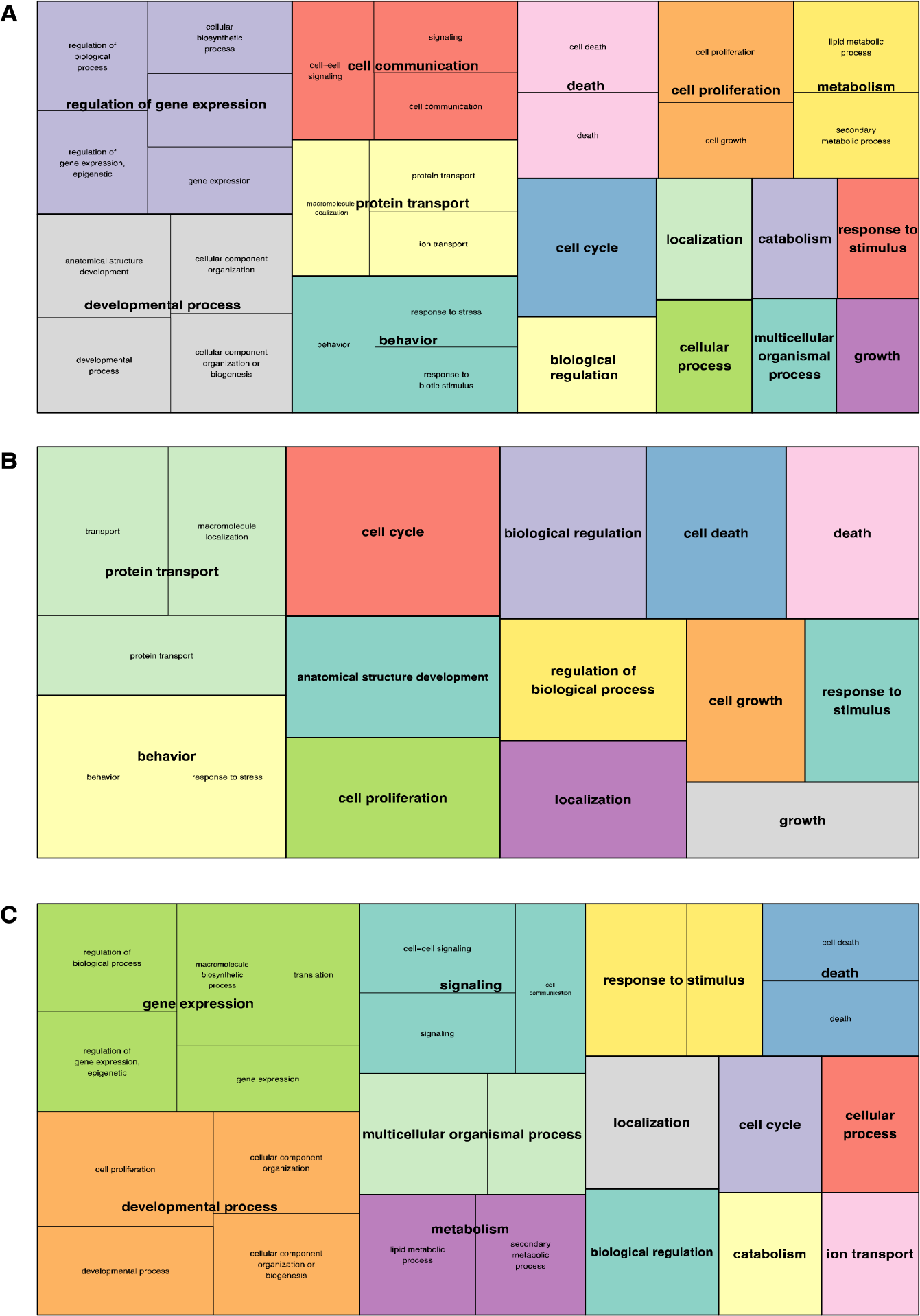
Gene Ontology Biological Process term enrichment for all growth and fecundity traits (A), adult *Daphnia* traits (B), and offspring traits (C). Figures are based on the reduction and summary provided by REVIGO (Supek et al. 2011), given the enrichment calculations from GOstats (Gentleman 2011). Large font labels are representative terms for the colored block over which the label is situated, and block size is proportional to the p-value of the enrichment (i.e., larger blocks possess lower p).

One mechanism driving gene coexpression is shared expression regulators, and by extension, shared recognition sequences in promoter regions. We identified between one and 17 promoter motifs enriched in the promoters of genes associated with each trait (SI Table 6). The number of motifs recovered per trait was proportional to the number of genes in each coexpression module considered, and support for each motif ranged from E-value < 1e^−3^ to 1.4e^−26^. Motifs containing TGC repeats, such as Motif 1 (MTGCTGCTGCTGCTGYY) of first-brood offspring mass “turquoise” module, were associated with half of all growth and fecundity traits, suggesting that it may be a target of a general transcriptional regulator associated with size and reproduction; there was no strong sequence similarity with known *Drosophila* motifs, however. Several motifs with long cytosine repeats, such as the motif with the lowest E-value (CYCCCCCCCCCCYHYHB, 1.43e^−26^), are highly similar to the *Drosophila* motif target of CG7368, an unnamed transcription factor associated with phagocytosis [65]. The *Daphnia* gene with the highest sequence similarity to CG7368, ZFP-ZMS1 (hxAUG26us24g213t1), was differentially expressed but not strongly associated with variation in any of the traits.

### An integrated network hypothesis

To generate a draft systems network hypothesis that spanned SNPs, DEGs, and traits, we intersected the DEGs strongly associated with SNP variation and trait variation to identify a systems genetic—i.e., spanning from genetic to expression to organismal variation—hypothesis for several growth and fecundity traits (Figure 7). This hypothesis is restricted to SNPs occurring in genes with transcription regulation annotations; although any of the SNPs associated with gene expression or trait variation (SI Table 7) may be causal or linked to causal variants, the annotations associated with these SNPs suggests a plausible regulatory mechanism. One of the markers is within an ecdysone-induced protein (hxAUG26us17g279t1) that is not differentially expressed, but the allelic variants strongly predict (*p* < 1e^−7^) the expression levels of 27 DEGs. The MAPK ERK-a and PA2G4 markers are both associated with cell proliferation through different pathways [64], and are linked to time to first brood through associations with four DEGs.

**Figure 7.**
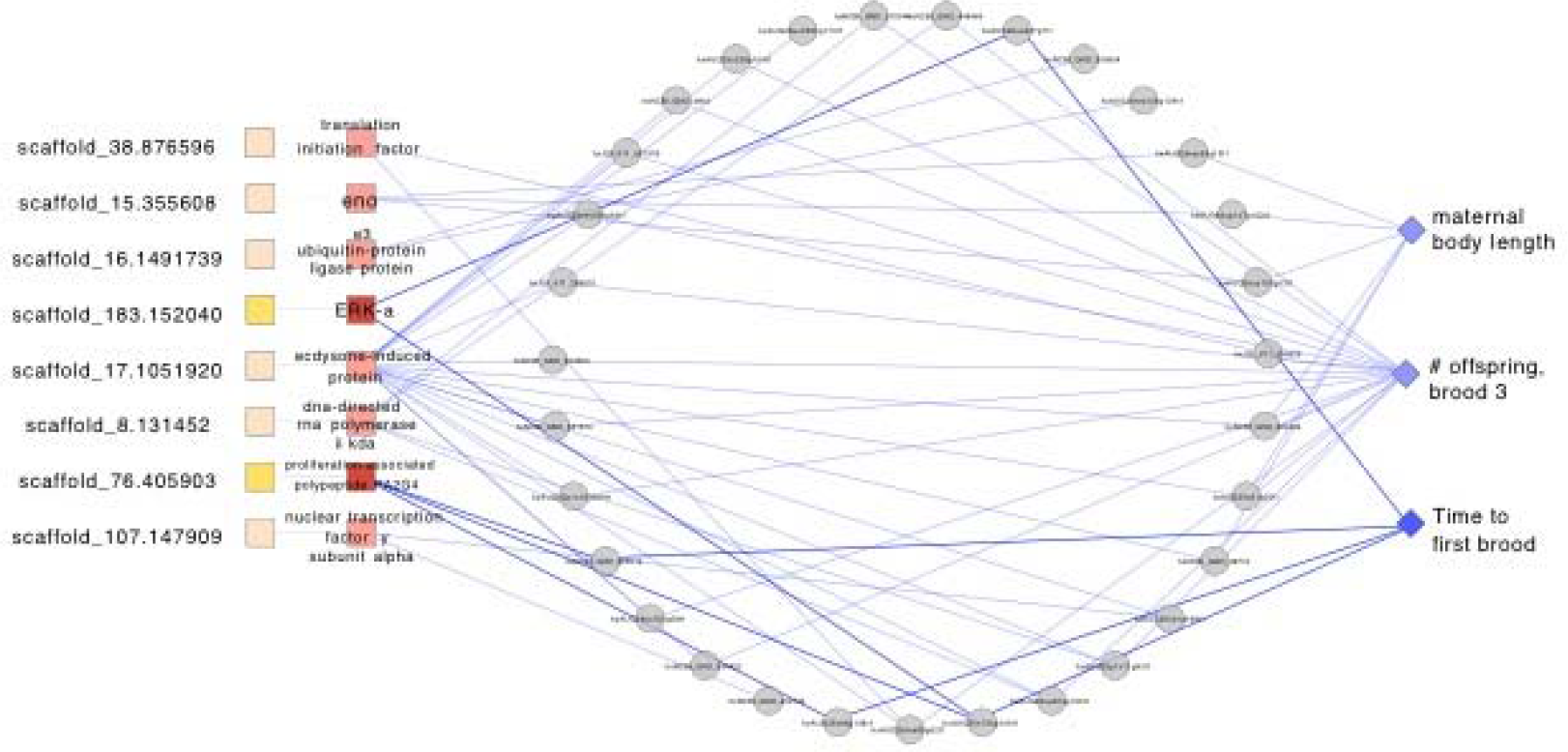
An integrated gene network hypothesis for three *Daphnia* life history traits. SNPs and the transcriptional regulators in which they are found are represented by squares; differentially expressed genes (DEGs) whose expression is related to marker variation and trait variation are represented by gray circles; and traits are represented by blue diamonds. Edges connecting marker-genes, DEGs, and time to first brood are highlighted darker blue. Labels and edges between coexpressed DEGs are suppressed for figure clarity; the data are available in SI Table X. Highlighted nodes and edges are discussed in the text.

## DISCUSSION

One goal of contemporary biology is to predict the drivers of variation between levels of organization, from genetic sequence to ecosystems. The importance of the core life history traits of size, growth, and fecundity in shaping ecological dynamics has long been hypothesized, is encompassed by ecological theory, and we have empirical support of their importance in a few systems, including the ecological model organism, *Daphnia pulex*. The recent sequencing of the *Daphnia* genome means that we can begin to relate genetic, trait, and ecological variation, as well as identify the genes underlying rapid evolution and its ecological consequences. In this paper we provide a first pass at the map from genetic to trait variation in traits known to affect *Daphnia* ecology, given a panel of wild-caught clones. We find substantial variation at three levels of biological organization—genotype, gene expression, and organismal phenotype—while linking the levels and generating novel hypotheses to test as the *Daphnia* system matures for ecological genomic research.

The amount of observed natural phenotypic variation was not surprising given the variety of habitats—abiotic, competitive, and predation regimes—from which the panel was collected. Previous studies of *Daphnia* that measured organismal traits also recovered substantial levels of variation (e.g., [35, 39, 65, 66]) among clones, but often were more focused on interspecific differences. Some trait correlations qualitatively matched those of previous studies; for example, Spitze and colleagues found a strong positive correlation between growth rate and reproductive output [35], as we found. However, while we found effectively no correlation between time to first brood and brood size, Spitze and colleagues recovered a negative correlation (−0.08 to −0.69, depending on the brood). We did not observe sign change between genetic and phenotypic correlations for any trait pair, and the degree of change was within expectations given heritabilities [68].

Our genome-wide genotyping offers one of the first population genomic examinations of the *D. pulex* group. While the panel is relatively small, we sampled individuals of all three ecotypes (two lake, fourteen shaded-pond, 16 sunny-pond) and found extensive genome-wide genetic variation with relatively little population structure. Coupled with the continuously distributed variation among the fourteen traits, this result supports reinforces other recent findings [69–71] suggesting continued gene flow between the ecologically isolated *D. pulex* and *D. pulicaria*, through hybrid back-crosses. Future population genomic studies with larger samples, and from across a larger geographic area, will shed light on the enigma that is the *D. pulex* group.

Our analysis recovered a highly structured and modular transcriptome in *Daphnia*. This finding reflects our expectations given the mechanisms of expression control (i.e., relatively few transcriptional regulators with many targets) and the results of similar studies [57, 72].

Enrichment for a relatively small number of Gene Ontology terms per module, and the presence of well-supported promoter motifs among coexpressed genes, further supports the cohesion of the modules. Putative functional annotations may be transferred to currently un-annotated DEGs (~ 66% of DEGs) given the patterns of coexpression and shared promoters with functionally annotated genes [73], advancing our knowledge of *Daphnia* molecular biology while generating hypotheses that can be tested by more focused approaches. Such extension is, however, beyond the scope of the present work.

In addition to support for several a priori expectations—from ecdysone-activated genes and gustatory receptor gene expression, to enrichment for a variety of biological terms—we also identified hundreds of candidate genes associated with variation in the fourteen *Daphnia* growth and fecundity traits. This magnitude of discovery is expected from genome-wide approaches and provides a foundation for novel hypothesis-driven molecular research with *Daphnia*. For example, the fact that >60% of DEGs associated with *Daphnia* growth and fecundity variation have no functional annotation provides fertile ground for new discoveries and insights into the functional diversification of genes. Other results are better-known: the central roles of *Partner of bursicon*, ERK-a, and Rab-32 to *Daphnia* size variation networks captures these genes’ known roles in development and cell proliferation in *Drosophila*.

The coexpression network analyses add to our collective knowledge about the inherent modularity of biological systems. Modularity—sets of interactions that are stronger within than between groups—is a common feature of biological systems, from neural networks to genetic architecture [74, 75]. Modularity itself may be a target of selection by decreasing interference between the modules [76], and intermediate levels of pleiotropy across modules should facilitate evolvability [77, 78]. Among the coexpression networks associated with variation in *Daphnia* traits examined here we found substantial variation in the degree and details of modularity. For example, although offspring and adult mass are the same character from different life stages, differences in the number of modules (three vs. five) and number of high betweenness-centrality genes (two vs. five) associated with each trait suggests different evolutionary potentials. How these coexpression modules are remodeled [79] across the developmental timeline, and in a variety of habitats, is an open question; the results provide a reference against which future research can be compared.

Variation in the GO terms enriched for the sets of genes associated with individual and groups of *Daphnia* growth and fecundity traits highlights similarities and differences in the underlying biological processes. Cell cycle, proliferation, and death are common between the groups of traits, but enrichment for protein transport and localization is restricted to adult size traits, while developmental processes are (as expected) enriched in offspring. We are not aware of similar analyses and comparisons in other species, but this observation suggests that quantifying numerous similar traits—rather than just one or two representatives of a trait class—has the potential to inform our understanding of the molecular basis of many small phenotypic distinctions [80, 81]. Note, however, that inferences from biological term analysis need to be tempered with the fact that most *Daphnia* genes are lineage-specific [11] and currently possess no functional annotation.

The goal of systems genetic approaches is to identify and understand the functional relationships between genetic variants; the genes whose expression (and post-transcriptional modifications) are affected by the genetic variants and environmental causes; and ultimately phenotypic variation [82, 83]. The combination of extensive genotype, expression, and phenotype data allowed us to generate a systems genetic hypothesis linking variation across these three levels of organization for *Daphnia* growth and fecundity. The relatively small size of the panel and lack of controlled crosses precluded a full probabilistic analysis of the network, and the focus in particular on genetic variants in “known” transcriptional regulators is cautious: for example, by requiring transitivity from genotype to organismal trait (i.e., markers associated with DEGs and one or more traits, with the DEGs also correlated with variation in the same trait) we restrict the core hypothesis (Figure 7) to eight markers and three traits. Many other loci and genes likely underlie growth and fecundity variation in *Daphnia*, but the elements of this network are supported in various analyses and provide hypotheses to be tested. For example, the systems hypothesis suggests a major role for an ecdysone-induced protein on scaffold 17; as a major regulator of arthropod molting [84], the role of ecdysone (or rather, the targets of ecdysone) is a strong candidate for general regulation of *Daphnia* growth and fecundity. Future work that perturbs the *Daphnia* system—through crosses, RNAi [85], plasmid integration [86], environmental manipulations [87, 88], and other methods (see, e.g., [89])—will help expand upon and test the present hypothesis.

While the present work is a distinct advance for *Daphnia* genomics, the limitations of the approach used here must be recognized. First, the expression and phenotypic data were collected in a single, benign common garden setting, and *Daphnia* is well-known for its extensive phenotypic plasticity (e.g., [89–92]). These results provide a baseline against which future work, under a variety of ecologically interesting and realistic conditions, can be compared. Second, we used whole *Daphnia* for RNA collection, but tissue-specific expression differences are a well-known phenomenon among many organisms and genes [93]. Future research that aims to isolate expression to particular *Daphnia* tissues will certainly refine our understanding of the relationship between genotype, expression, and morphological variation.

There are two main implications of the present research for our collective understanding of *Daphnia* ecology and evolution. First, this is one of the few examples, if not the only example, of ecological genomics [94, 95] applied to an organism whose community ecological context is very well-studied. Because the implications for size and fecundity variation in *Daphnia* are known to extend both up and down trophic levels, identifying loci and genes affecting these traits drives at causes of the extended phenotype [96]. That is, if the genotype at scaffold_17:1051920 (a T/C or T polymorphism) affects *Daphnia* size (through numerous intermediary genes), then it is therefore predictive of the effects on phytoplankton communities through grazing pressure, and susceptibility to predation by vertebrate and invertebrate predators. This example and the hundreds of others discovered in this work provide numerous hypotheses to be tested as explanations of organismal and community variation.

Second, it has become apparent over the past decade that rapid evolution—changes occurring on the scale of just a few generations—can have dramatic ecological implications [15, 97–99]. *Daphnia* has been a model system for examining eco-evolutionary dynamics of disease [100], predation [101, 102], and eutrophication [14], and in each of these cases the basic life history traits studied here play a central role in shaping the ecological interactions. An outstanding issue is identifying the loci underlying evolutionary change of ecological importance. For example, phenotype data alone cannot answer whether the same loci are involved in parallel bouts of adaptation, or if there are multiple, unique avenues of adaptation (see, e.g., [103, 104]). Because the panel was collected from natural populations, the results provide candidates for the loci that may be responsible for rapid, ecologically important evolution in *Daphnia*.

## CONCLUSION

Here we have identified genetic and gene expression variants associated with variation in life history traits of the model ecological organism, *Daphnia pulex*. In addition to recovering several expected gene/pathway relationships, the analyses uncovered numerous novel gene-trait relationships that form the basis for future hypothesis-driven research. These results are an important first step for understanding the molecular basis of variation in *Daphnia* growth and fecundity—traits for which the impacts of variation extend across communities—and are candidates for the molecular basis of ecologically important adaptation. Furthermore, these results may be vital resources in comparative analysis of the molecular evolution of organismal variation across the tree of life.

## METHODS

### Clone collections and maintenance

Nominal *Daphnia pulex*, *D. pulicaria*, and hybrid clones were collected from a variety of waterbodies in the area surrounding Kellogg Biological Station, Michigan, USA, in June 2009. The panel includes clones from the mesocosm experiments of [105]. Individuals were isolated and cultures started in Austin, TX, from a single female; after initial mortality of isolates and clone losses during the assay period, we obtained 32 unique lineages. All working cultures were maintained in ADaM media [106] in an environmental chamber at 20**°**C with 16:8 L:D cycles. They were fed daily 1-2ml Shellfish Diet (Reed Mariculture, Campbell, California), at 2×10^6^ cells per milliliter, regularly supplemented with live *Scenedesmus acutus*.

### Phenotypic assays

*Daphnia* are model organisms for studying maternal effects [107, 108], but such effects are not the focus of this work. To minimize maternal and grand-maternal effects on size, growth, and fecundity estimates, each replicate was reared through two generations of single-individual breeding before assaying the focal generation. All individuals in the assays were raised in 100ml cups in ADaM media and fed 1ml shellfish diet each day, supplemented with 4×105 cells of *S. acutus* every-other-day. Half of the media was replaced in each cup every 3d, and each cup was thoroughly cleaned and all media replaced each week. Each cup was checked daily for the presence of offspring and for mortality. If offspring were present, then they were counted and 2-4 placed into a new cup with fresh media and food. These individuals were randomly culled to a single individual within 2d of their birth, but young individual mortality required that >1 be retained initially. If the focal individual had died at the check, then the replicate was started over at the grand-maternal generation with an individual taken from the working culture.

Once the focal generation had been reached, we recorded the dates and sizes of the first, second, and third broods. All offspring of the first and third broods were collected and stored in a 1.5ml tissue tube in 95% ethanol in a −10°C freezer. The mother was collected and stored in the same manner at the release of her third brood. At regular intervals, we removed samples from storage for further measurement. Adult body length to the base of the tail spine, body depth at the deepest point, and tail spine length were measured to the nearest 0.1mm under a 10-40x dissecting microscope. Adults and 1-15 juveniles were then placed in pre-tared aluminum mini weigh boats and dried for 48-72h at 60**°**C prior to weighing. *Daphnia* mass was measured to the nearest 0.1μg on a Sartorius ultramicrobalance placed in a closed room on a stabilizing marble bench. (Note that we rounded all measurements to the nearest 1μg for analysis.) Growth rate was calculated as the difference between log(mean neonate mass) and log(adult mass) divided by the number of days between the birth and the third brood. Adults were collected after weighing, and stored at −10**°**C for stoichiometric analysis. Percent phosphorus of each adult was established by measuring absorbance spectrophotometrically at 850nm.

Given the phenotypic data for three replicates of each clone, we calculated quantitative genetic parameters using a random effects model of form trait ~ 1 + (1 | clone) with the lme4 package [109] for R 2.15 [110]. Genetic correlations between traits was calculated from genotypic trait values and phenotypic correlations from all data.

### Genomic samples

We measured constitutive gene expression for each clone as a starting point for interrogating the genotype-phenotype map in *Daphnia*. To do so, we raised three replicate sets of cultures of each clone for RNA sampling in the same environmental chamber, and using the same feeding regimen, as the working cultures. Each culture was maintained at a low density of 8-12 individuals per 150ml ADaM, for three generations. Even at this low density and high food provision, some clones appeared to be producing ephippia (resting stage eggs) whereas other clones were not. Collections were marked if there was any sign of ephippia production, which would likely alter gene expression profiles. Three individuals were collected from each replicate and each was stored in individual collection tubes to facilitate single-individual analysis of expression. That is, we collected a total nine individuals per clone. The collections took place on two adjacent days from 10:00-13:00h local time to minimize any circadian effects. Samples were placed immediately into a liquid nitrogen-filled Dewar, then transferred to and stored in a −80°C freezer.

After grinding by mortar and pestle in a liquid nitrogen bath, RNA was extracted from single individuals using Qiagen RNeasy kits (Qiagen, CA) per the manufacturer’s instructions. RNA preparation for SOLiD sequencing followed the basic method of Meyer and colleagues [111]. In brief, this is a 3’ tag RNA-Seq method whereby fragmented RNA is reverse-transcribed to a cDNA library, amplified, and tagged with a SOLiD-ready barcode. Initial fragmentation was accomplished by a single 3-minute period at 95°C in a thermocycler; fragmentation was confirmed by gel analysis. After fragmentation, a cDNA library was created by reverse transcription using SuperScript II reverse transcriptase and a switching template primer. We amplified the cDNA library using Titanium Taq and a thermocycle regimen of 5-min at 95C then 19 cycles at 95C (40s), 63C (1min), and 72C (1min). The PCR products were purified using a NucleoMag 96 cleanup kit, per manufacturer’s instructions. We quantified DNA concentrations after cleanup using a NanoDrop spectrophotometer. Barcodes were ligated to each sample using the SOLiD multiplex P1 oligo, 1uM barcode oligo, Titanium Taq, and a amplification profile of four cycles at 95**°**C (40s), 63**°**C (1min), and 72**°**C (1min). After amplification, samples were run on a 1x TBE gel, and size-selected between 180-250bp using a low molecular weight ladder (NEB #N3233S). The cDNA in each gel slice was extracted by immersing the gel slice in nuclease-free water overnight at 4**°**C. The 96 prepped libraries were given to the University of Texas Genome Sequencing and Analysis Facility (UT GSAF) for sequencing on SOLiD 5500XL and v4 platforms, with a target of 2 million 50-bp reads per sample.

Single *Daphnia* individuals did not yield sufficient gDNA for RAD genotyping, so we pooled three individuals of each clone for extraction. Genomic DNA (gDNA) was extracted using Qiagen DNeasy extraction kits (Qiagen, CA) per manufacturer’s instructions, with two exceptions: we did not vortex samples, in order to ensure that gDNA remained intact before *AlfI* digestion, and we completed the final rinse with 40μl nuclease-free water rather than 100μl to increase yield concentration. In brief, the 2b-RAD genotyping method (Wang et al. 2012) uses the *AlfI* restriction enzyme to digest the gDNA: SOLiD-system adaptors were ligated to the digested DNA and unique barcodes are then incorporated with the ligated products for each sample. The target constructs are 136bp in length and were extracted from electrophoretic gels. Samples of the 32 clones were prepared and sent to the UT GSAF for sequencing at a target of 1 million 50bp reads per sample.

### Genotyping

We used the GATK [112, 113] two-phase genotyping process to call genotypes from the filtered SAM files. First, we used the IndelRealigner to correct for insertion and deletion errors inferred during mapping, then performed an initial genotyping pass with UnifiedGenotyper (using – ploidy 2, -glm BOTH, and -out_mode EMIT_ALL_CONFIDENT_SITES). From the first-pass genotyping we extracted variant sites in the 90th percentile of both genotype qualities (GQ) and QUAL scores as high-quality genotypes to be used in base quality score recalibration. After applying BaseRecalibrator we performed a second round of genotyping and extracted variants with GQ > 30 (*p* < 0.001). Subsequent analyses based on genotypic variation used a subset of loci in which ≥ 78% of clones (≥ 25) were genotyped.

Two risks of using RNA-Seq derived reads for genotyping include allele-specific expression [114] and RNA editing [115]. To test if these potential sources of error were introducing significant bias, we genotyped all clones with DNA- and RNA-only data sources and compared genotype calls at overlapping sites: if either error source is prevalent, we expect to detect many more heterozygous loci from DNA-derived sequence. There were an average of 7122 overlapping loci per clone, and 10.3 differences on average, but both DNA- and RNA-derived genotypes possessed heterozygous calls when the alternate call was homozygous. Given the very low error rate (0.14%) and the fact that *Daphnia* genomics is a very young field, we opted to retain the joint genotype data with a very slightly higher false discovery risk.

### Expression analysis

We used the classic SHRiMP output and the probcalc function for expression analysis, classifying mappings with normodds > 0.66 (odds-ratio ≥ 2 for the next-best mapping) as uniquely mapped. Ambiguously mapped reads were allocated to multiple genes in proportion to the support for the mapping and the proportion of uniquely mapped reads assigned to each gene. While allocation is required to reduce bias against multi-copy versus single-copy genes, it is known that allocating multimapped reads biases low-expressed genes high [116], however, because low-coverage genes (< 5 reads per million mapped) were excluded from analysis this source of bias is negligible in the expression data. Last, to achieve comparability between clones with different sequencing coverage, all expression levels were converted to reads per million mapped and rounded to the nearest integer.

We used a generalized linear model (GLM) with quasipoisson errors, log link function, and variance inflation estimated for each gene from the pooled data to test for differentially expressed genes (DEGs), with clone ID as the predictor [117]. P-values were derived by likelihood ratio test against the null model, significance was set at p < 0.05, and Benjamini-Hochberg false discovery rate (FDR) control was exerted at a 1% rate.

### Marker- and gene-trait associations

We tested for marker-trait variation using a linear model and set significance at a nominal p-value of 1e^−5^. Only markers with at least three clones possessing the minor allele (i.e., 9-12% minor allele frequency, depending on the number of clones typed at a locus) were used in this analysis.

We used distance correlation, which is not dependent on linear or monotonic relationships between variables [118–120], to quantify the strength of association between organismal trait means and gene expression means. The p-values of distance correlations were derived from extensive bootstrapping of expression data, and FDR control exerted at the 1% level. Genes whose expression was significantly associated with variation in each trait were retained as initial candidates underlying variation.

### Coexpression networks and candidate gene refinement

We used WGCNA [121] to (a) identify gene coexpression modules associated with each trait and (b) refine the list of candidate genes. We applied WGCNA to the residuals of trait:gene regressions to reduce the occurrence of false-positives; three regression models (linear, log-limited, and negative exponential) were tested for each trait-gene combination because non-linear relationships from distance correlation were detected, and the residuals from the model with the highest R^2^ were used. Genes that did not cluster into a module (i.e., assigned the “gray” module of WGCNA) were removed from the candidate list as likely false-positives [57]. Modules were identified by estimating the exponent required for a scale-free distribution, and module membership refined by adjusting reassignThreshold and mergeCutHeight to best-match the modules apparent in the heat map. Additional flags included pamStage=FALSE, TOMType=”unsigned”, TOMDenom=”mean”, and minModuleSize=8. Network figures were created in Cytoscape by exporting the WGCNA data for the best-connected.

After identifying the final candidate gene list for each trait we used the GOstats package for R [122] to test for Gene Ontology (GO) term enrichment. We defined gene universes for (a) all *Daphnia* genes with GO annotations and (b) *Daphnia* DEGs with functional annotations. Because relatively few *Daphnia* genes possess functional annotation (~25%), we set pvalueCutoff at 0.1. Enrichment was quantified both at the level of all candidate genes for a trait and module-wise for the WGCNA-defined modules associated with variation in a trait. We used REVIGO [123] for GO term enrichment visualization.

We used XXmotif [124] to identify candidate promoter motifs shared among the genes and modules associated with variation in each trait. Promoters were extracted from the *Daphnia* genome file from 1000bp upstream and 200bp downstream of each gene’s transcription start site, both strands were searched for motifs, a background model of order 2 was employed, and a medium threshold was used for merging similar motifs. We retained motifs with E < 0.1, which corresponds to E < 0.01 because XXmotif estimates are biased high by approximately an order of magnitude for these sample sizes (see Supp. Info. 1 of [124]). We searched TOMTOM [125] for known motifs in *Drosophila* similar to those recovered with XXmotif.

To identify candidate regulators of trait-specific coexpression modules, we mined (case-insensitive grep) module members with annotations for “transcriptional regulation”, “kinase” and “MAPK”, because the known role of TFs, kinases, and particularly mitogen-activated phosphate kinases in transcription regulation. The integrated network of Figure 1.7 was created by intersecting the (transcription-associated) marker-DEG, DEG-DEG, and DEG-trait data to establish relationships across levels of biological organization.

## ACKNOWLEDGEMENTS

This work was supported by grants from the University of Texas at Austin (UT) to JWM and MAL and NSF DEB 0717370 to MAL. Sequencing was completed by the UT Genome Sequencing and Analysis Facility, and the Texas Advanced Computing Center provided access to the Lonestar supercomputing resource for many of the analyses reported here. We thank Mike Pfrender and Misha Matz for providing helpful comments on earlier drafts.

